# Breaching the diatom frustule: *Alteromonas macleodii* triggers protoplast emergence and programmed necrotic-like cell death in the oceanic diatom *Thalassiosira rotula*

**DOI:** 10.1101/2025.11.20.688403

**Authors:** Isobel Short, Clara Martínez-Pérez, Roman Stocker, Uria Alcolombri

## Abstract

Diatom-bacteria interactions drive biogeochemical fluxes at the base of the marine food-web. While the role of bacteria in recycling organic matter is well established, their potential to actively induce algal death remains poorly understood. Here we examined the interaction between the bloom-forming diatom *Thalassiosira rotula* and a known member of its microbiome, *Alteromonas macleodii*. Microscopy revealed that *A. macleodii* breached the diatom’s silica frustule, inducing >95% of diatom cells to emerge from their shells within 72 hours, exposing protoplasts to lysis and exploitation. This algicidal interaction is species-specific and contact-dependent. Dual transcriptomics revealed programmed necrotic-like cell death underpins *T. rotula*’s terminal morphological change, alongside bacterial upregulation of motility, chemotaxis, and coordinated metabolism to exploit the diatom host. These findings indicate the presence of a programmed necrosis-like pathway in diatoms, akin to that known in mammals. They also reveal a novel algicidal strategy, recasting *A. macleodii* as a specialized antagonist capable of influencing bloom termination and removing the ballast provided by diatom frustules, thereby reshaping carbon cycling in the ocean.

## Introduction

Marine microbial ecosystems are composed of networks of interactions among species, which occur at the microscale yet influence biogeochemical cycles at global scales. Having cohabited the surface oceans for millions of years, diatoms and bacteria have co-evolved a strong ecological coupling^1^. Diatom-bacteria relationships exist on a continuum from reciprocal to antagonistic interactions, encompassing mutualism, commensalism, parasitism and algicidal interactions^1,2^. Interactions with bacterial partners influence diatom metabolism^3^, productivity^4^, cell morphology^5^, aggregation^6^, cell division^7^, and sexual reproduction^8^. By shaping the metabolism and life cycle of their algal protagonists, these relationships affect productivity, microbial community dynamics and oceanic carbon cycling. Diatoms’ siliceous frustules ballast organic particles, enhancing carbon flux to depth and making diatom-derived a major contributor to particulate organic carbon (POC) export^9^. While bacteria-driven aggregation can modulate carbon export^6^, interactions that disrupt this ballast may have similarly significant consequences, an effect that remains unexplored.

Bacteria most commonly associated with diatoms belong to only a few genera within the phyla Proteobacteria and Bacteroidetes^10^, highlighting the extensive specialisation required to establish stable interkingdom interactions. For example, to achieve favourable rates and durations of encounters with algae, bacteria have evolved targeted chemotaxis and attachment mechanisms^11–14^. Once the two actors are in close proximity, they are able to more effectively exchange the chemical currencies mediating their interaction, including micronutrients such as vitamins^15^ and iron^16^, along with nitrogen^17^ and dissolved organic carbon (DOC)^18,19^. These resource exchanges occur primarily within the phycosphere — the micro-environment enriched in algal-derived metabolites surrounding individual phytoplankton cells^2,20^ — where they sustain individual partners and collectively shape the structure, productivity, and biogeochemical impact of marine microbial communities.

In the most extreme form of antagonism, some heterotrophic bacteria have evolved to secure nutrients by actively inducing phytoplankton cell death^21–23^. These algicidal strains deploy diverse mechanisms, inducing growth inhibition^7^, morphological changes^24^, and most dramatically, cell lysis^25^, processes that modulate microbial community dynamics and profoundly impact the balance between primary and secondary production in marine ecosystems^26^. Similar to viruses, which regulate microbial ecosystems by driving phytoplankton mortality and fueling the viral shunt^27,28^, algicidal bacteria can disrupt bloom dynamics^21–23,25^, yet their ecological roles, prevalence, and antagonistic mechanisms remain poorly characterized. Moreover, our understanding of the cellular basis of algal mortality^29^ lags behind that of other eukaryotes, such as mammals and yeast, where distinct cell death pathways are well defined with regards to their molecular hallmarks^30,31^. Given their differing consequences for nutrient release to the surrounding environment, resolving these fundamentals of phytoplankton cell death is essential for understanding their impacts on microbial ecology and nutrient fluxes in the ocean.

Here, we report the identification and characterization of a new form of algicidal interaction between two globally dominant microplankton species, the diatom *Thalassiosira rotula*^32,33^ and the bacterium *Alteromonas macleodii*^34,35^. *A. macleodii* is emerging as a model marine heterotroph, having evolved versatile copiotrophic strategies to thrive in nutrient-rich niches^36–43^, and is also a core member of the microbiome associated with *T. rotula*^44^. Using time lapse microscopy, we observed that co-culture with the bacterium induces a terminal morphological change in the diatom, causing complete emergence of the protoplast from the silica shell and frequently leading to cell lysis. *A. macleodii* exhibits rapid population growth as a result of the interaction. This response is species-specific among all those we tested and contact-dependent. Employing a dual transcriptomic approach allowed us to follow the simultaneous gene expression changes in both species during co-culture, revealing that *A. macleodii* induces a stress response in *T. rotula* that leads to a programmed cell death (PCD) pathway, with genetic and phenotypic signatures of programmed necrosis. Bacterial gene expression analysis revealed that motility, chemotaxis and dynamic metabolic strategies underlie *A. macleodii*’s virulence and its ability to exploit diatom exudates and lysates, highlighting that traits previously considered opportunistic can also serve algicidal functions. Our results reveal a novel algicidal mechanism that shapes phytoplankton and bacterial population dynamics and, by removing the ballast provided by diatom frustules, may alter the sinking speed of particles and fate of carbon in the ocean.

## Results

### *A. macleodii* triggers protoplast formation in *T. rotula*

To investigate their interaction, exponentially growing *T. rotula* CCMP 3096 was co-cultured with *A. macleodii* ATCC 27126 under controlled light and temperature conditions (Methods). *A. macleodii* was added at a final concentration of 1 × 10⁶ cells/mL from exponential-phase cultures grown in 2216 MB, resulting in 1% (v/v) 2216 MB in co-cultures, a source of DOC. After 24 h of co-culture, a high proportion of *T. rotula* cells were observed to have undergone a striking morphological change: in each case, the cylindrical cell bent at the midpoint as a membrane protrusion formed at the thecae junction (Fig. 1a), and expanded within hours until the cell fully emerged from its silica frustule, resulting in a spherical protoplast (Fig. 1a, b, Supplementary Movie 1). Protoplasts retained intact nuclei and chloroplasts (Fig. 1c). Within 72 h of co-culture, over 95% of the diatom cells showed this transformation, significantly more than the 3% in control cultures without *A. macleodii* (one-way ANOVA with Tukey’s post-hoc test, *P* < 0.001; Fig. 1d). This effect was initially observed in xenic *T. rotula* with their natural microbiome. We then repeated the co-culture experiments with axenic *T. rotula*, confirming a high proportion of protoplasts after 72 h and demonstrating that the effect was independent of the presence of other bacteria (Fig. 1d). Addition of bacterial culture medium (1% 2216 MB) alone induced significantly fewer protoplasts than addition of *A. macleodii*: 10% in xenic cultures (significantly above the no-treatment control; *P* < 0.001) and 3% in axenic cultures (not significantly different than the no-treatment control; *P* > 0.05). The increase in protoplasts upon 2216 MB addition in xenic but not axenic cultures possibly reflects DOC-mediated activation of naturally occurring *A. macleodii* in the xenic cultures, where *A. macleodii* was detected by 16S rRNA analysis (Supplementary Table S1). These results suggest that, in the presence of low concentrations of DOC, *A. macleodii* induces the dramatic morphological transformation in *T. rotula*.

**Figure 1.**
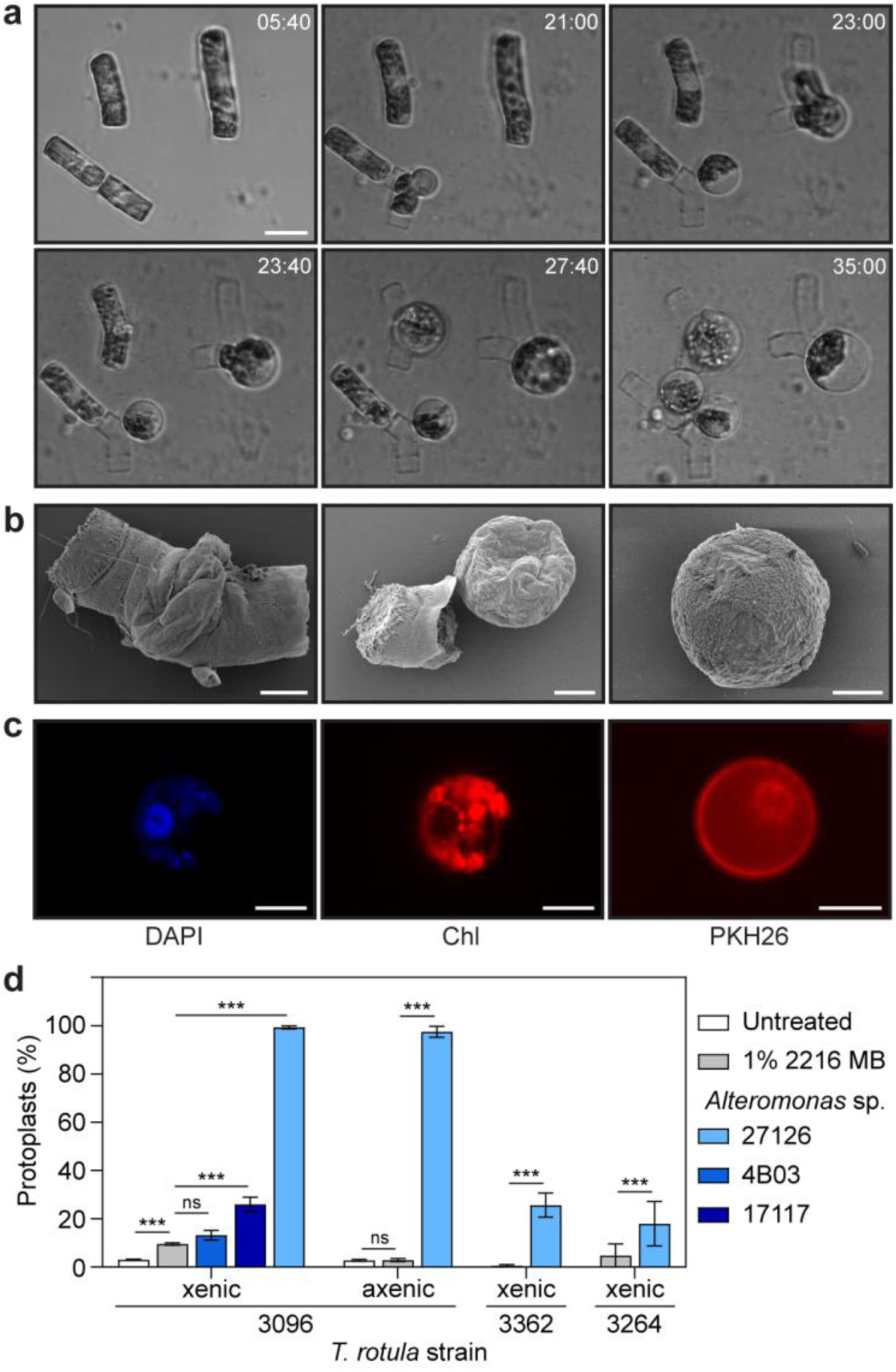
*A. macleodii* induces *T. rotula* to emerge from its protective silica frustule. **a** Representative images from a time-lapse video of four *T. rotula* CCMP 3096 cells emerging from their silica frustules and forming protoplasts, in co-culture with *A. macleodii* ATCC 27126. Labels indicate time since the start of co-culture (hh:mm), scale bar 20 μm (full video in Supplementary Movie 1). **b** Scanning electron micrographs of co-cultures showing progressive formation of *T. rotula* protoplasts during co-culture. Left: membrane protrusion emerges between the thecae. Centre: protoplast released from the frustule. Right: detached protoplast. Scale bars 5 μm. **c** Fluorescence microscopy images of *T. rotula* protoplasts showing cellular arrangement. Left: DAPI staining of DNA. Centre: chlorophyll autofluorescence of the chloroplasts. Right: PKH26 staining of the cell membrane. Scale bars 10 μm. **d** Proportion of protoplasts (% of total diatom cells) in three *T. rotula* strains (with different microbiomes) after 72 h of exposure to one of three bacterial strains, fresh bacterial culture medium (1% 2216 MB), or left untreated. Bars show means ± s.d. (*n* = 3 independent biological replicates), >100 diatom cells examined per replicate by phase contrast microscopy after 72 h co-culture. Horizontal bars show statistical significance from one-way ANOVA with Tukey’s post-hoc test; *** = *p* < 0.001, ns = not significant (*p* > 0.05).

The antagonistic interaction between *A. macleodii* and *T. rotula* is highly specific. We observed no morphological changes in *T. rotula* CCMP 3096 co-cultured with two other algal-associated bacteria, *Ruegeria pomeroyi* DSS-3 and *Vibrio alginolyticus*. Likewise, *A. macleodii* ATCC 27126 caused no morphological changes in the diatoms *Thalassiosira weissflogii* CCMP 1050, *Thalassiosira pseudonana* CCMP 3367 or *Phaeodactylum tricornutum* CCMP 2560. We observed no significant increase in protoplast formation in *T. rotula* CCMP 3096 co-cultured with *Alteromonas* sp. ALT199 strain 4B03 (*P* > 0.05; Fig. 1d). Meanwhile, *Alteromonas mediterranea* DSM 17117, a close relative, induced protoplast formation in 26% of *T. rotula* CCMP 3096 cells, significantly higher than the 1% 2216 MB control (*P* < 0.001), but much lower than the 94% induced by *A. macleodii* ATCC 27126 (*P* < 0.001). Though less strongly than *T. rotula* CCMP 3096, strains CCMP 3264 and 3362 also showed a significant increase in protoplast formation upon co-culture with *A. macleodii* ATCC 27126 compared to the 2216 MB control, averaging 18% and 25% of cells forming protoplasts after 72 h, respectively (*P* < 0.001, versus 2216 MB control for both strains). These results suggest that the interaction we have described is likely species-specific, occurring only between certain *Alteromonas* species and *T. rotula* strains, with the strongest effect among the pairs that we studied occurring between *A. macleodii* ATCC 27126 and *T. rotula* CCMP 3096, which were therefore used for all subsequent experiments.

### Contact dependent algicidal activity of *A. macleodii* against *T. rotula*

Time-lapse microscopy and flow cytometry revealed three distinct phases in the *T. rotula* - *A. macleodii* interaction (marked in Fig. 2a, b, f). Phase 1 (0-10 h) involved growth arrest in *T. rotula* (Fig. 2a). In Phase 2 (10-35 h), *A. macleodii* reached maximum growth (doubling time 5.9 ± 1.1 h; Fig. 2a) as diatom cells rapidly transformed into protoplasts, with >50% of diatom cells evacuating their silica shell by 20 h and 95% after 35 h (Fig. 2b). This transformation was irreversible: we never observed protoplasts regaining the cylindrical morphology or forming new frustules. Instead, protoplasts frequently underwent lysis, starting from 35 h after initiation of co-culture and defining Phase 3. By 72 h of co-culture, when the experiment ended, 27% of the protoplast diatom cells had lysed (Fig. 2b).

**Figure 2.**
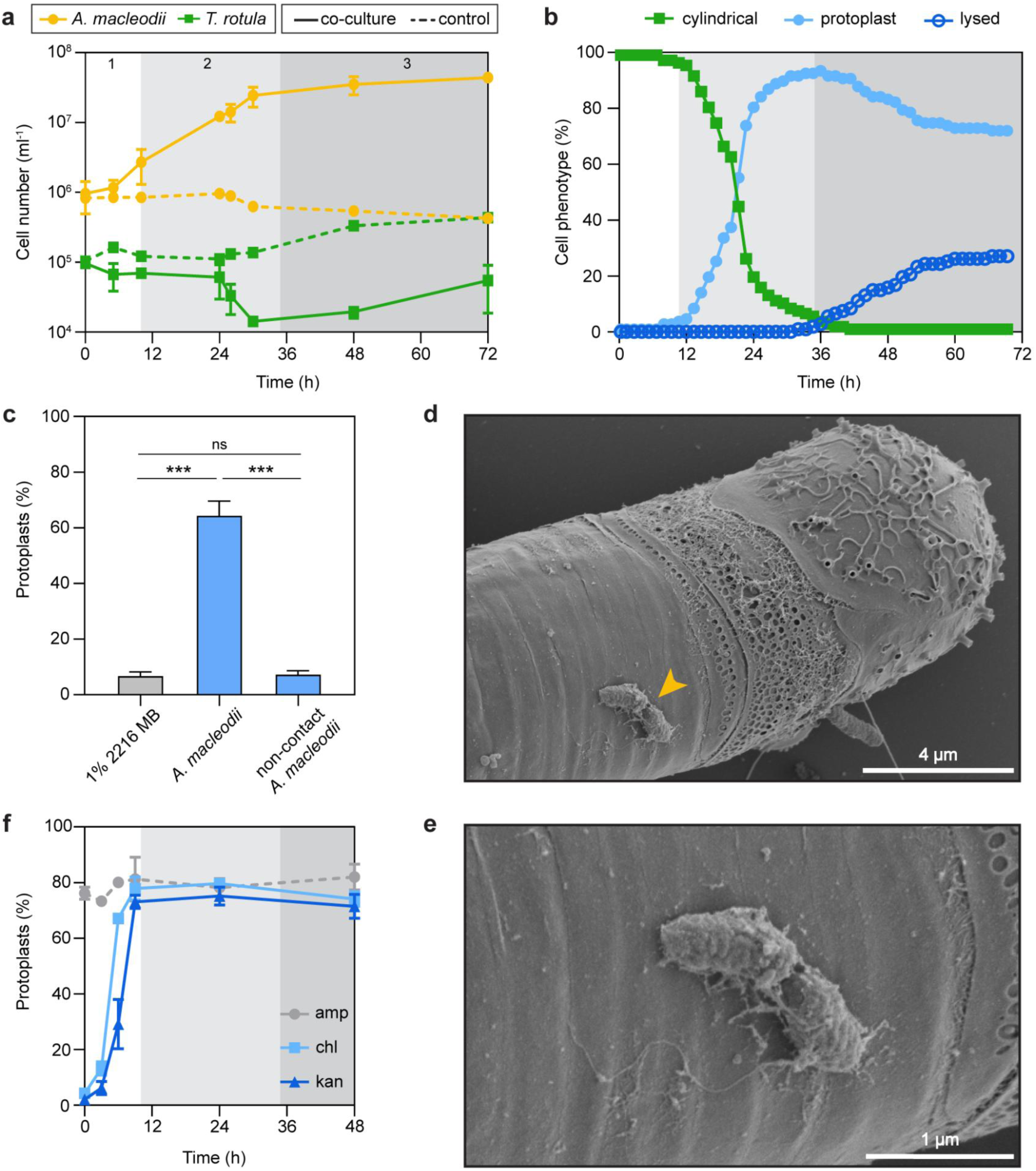
The antagonistic interaction between *T. rotula* and *A. macleodii* requires direct contact and occurs rapidly and irreversibly. **a** Population size over time for *A. macleodii* and *T. rotula* in co-culture and when cultured alone (“control”). Symbols represent mean ± s.d. (n = 3 independent biological replicates). Shaded regions denote the three phases of the interaction: Phase 1 (0–10 h) is associated with growth arrest in *T. rotula*; Phase 2 (8–35 h) coincides with the maximum growth rate of *A. macleodii*; and Phase 3 (35–72 h) involves lysis of *T. rotula* cells and is described in subsequent panels. **b** Proportion of *T. rotula* cells exhibiting cylindrical, protoplast or lysed phenotypes over time during co-culture with *A. macleodii*, as determined by video microscopy tracking of >100 individual *T. rotula* cells over the 72 h co-culture period. *T. rotula* cells undergo the morphological transition during Phase 2 and lysis begins in Phase 3. **c** Protoplast formation (% of diatom cells) in *T. rotula* after 72 h of culture with 1% 2216 MB (negative control), in co-culture with *A. macleodii*, or in co-culture with *A. macleodii* in shared medium with physical contact prevented by a filter (“non-contact *A. macleodii*”; Methods). Bars show mean ± s.d. (n = 3 independent biological replicates), >100 cells counted per replicate. Horizontal bars indicate statistical significance from one-way ANOVA with Tukey’s post-hoc test; *** = p < 0.001, ns = not significant (p > 0.05). **d** Scanning electron micrograph of a co-culture fixed after 5 h, showing *A. macleodii* on the surface of a *T. rotula* cell. Scale bar 4 µm. **e** Magnified view of D. Scale bar 1 µm. **f** Protoplast formation (% of diatom cells) in *T. rotula* after 72 h of co-culture with *A. macleodii*, following the addition of ampicillin (amp), chloramphenicol (chl), or kanamycin (kan) after 0, 3, 6, 9, 24 or 48 h. Symbols show mean ± s.d. (n = 3 independent biological replicates), >100 diatom cells counted per replicate.

The antagonistic interaction required direct contact. No significant increase in the prevalence of protoplasts was observed in *T. rotula* when the two organisms were separated by a 0.22 μm membrane (“non-contact”), compared to the 2216 MB negative control (one-way ANOVA with Tukey’s post-hoc test, *P* < 0.001; Fig. 2c). Furthermore, scanning electron microscopy (SEM) showed bacteria attached to diatom cells within Phase 1, after only 5 h of co-culture (Fig. 2d, e).

*T. rotula*’s fate is determined within 9 h of exposure to *A. macleodii*. Antibiotic inhibition of protein synthesis in *A. macleodii* with chloramphenicol (“chl”) or kanamycin (“kan”) showed that commitment to the morphological change in *T. rotula* occurs during Phase 1 - before protoplast formation in Phase 2 - and is irreversible even if *A. macleodii* is subsequently inhibited. While antibiotic inhibition before 9 h produced a marked reduction in the proportion of protoplasts at 72 h relative to the negative control ampicillin (“amp”, to which *A. macleodii* is resistant), addition at or after 9 h yielded values comparable with the control (Fig. 2f).

### Protoplast formation in *T. rotula* is linked to programmed necrotic-like cell death

To understand the responses of *T. rotula* and *A. macleodii* at the level of gene expression during their interaction, we conducted dual transcriptomic sequencing of co-cultures and controls at 8 time points over 72 h (0, 5, 10, 24, 26, 30, 48, 72 h), with 5 biological replicates per time point. The *de novo* assembled transcriptome of *T. rotula* contained 35,541 unique predicted coding sequences (CDS), of which 14,335 genes (40%) were differentially expressed in co-culture relative to the negative controls (likelihood ratio test, adjusted *P* < 0.1; Methods). Multivariate analysis showed samples clustered by timepoint, indicating a reproducible progression of the transcriptomic response (Fig. 3a): during the first 10 h (Phase 1), gene expression profiles were similar in co-cultures and controls, while they diverged markedly from 24 h onwards (Phase 2), coinciding with the phenotypic change in the diatom population.

**Figure 3.**
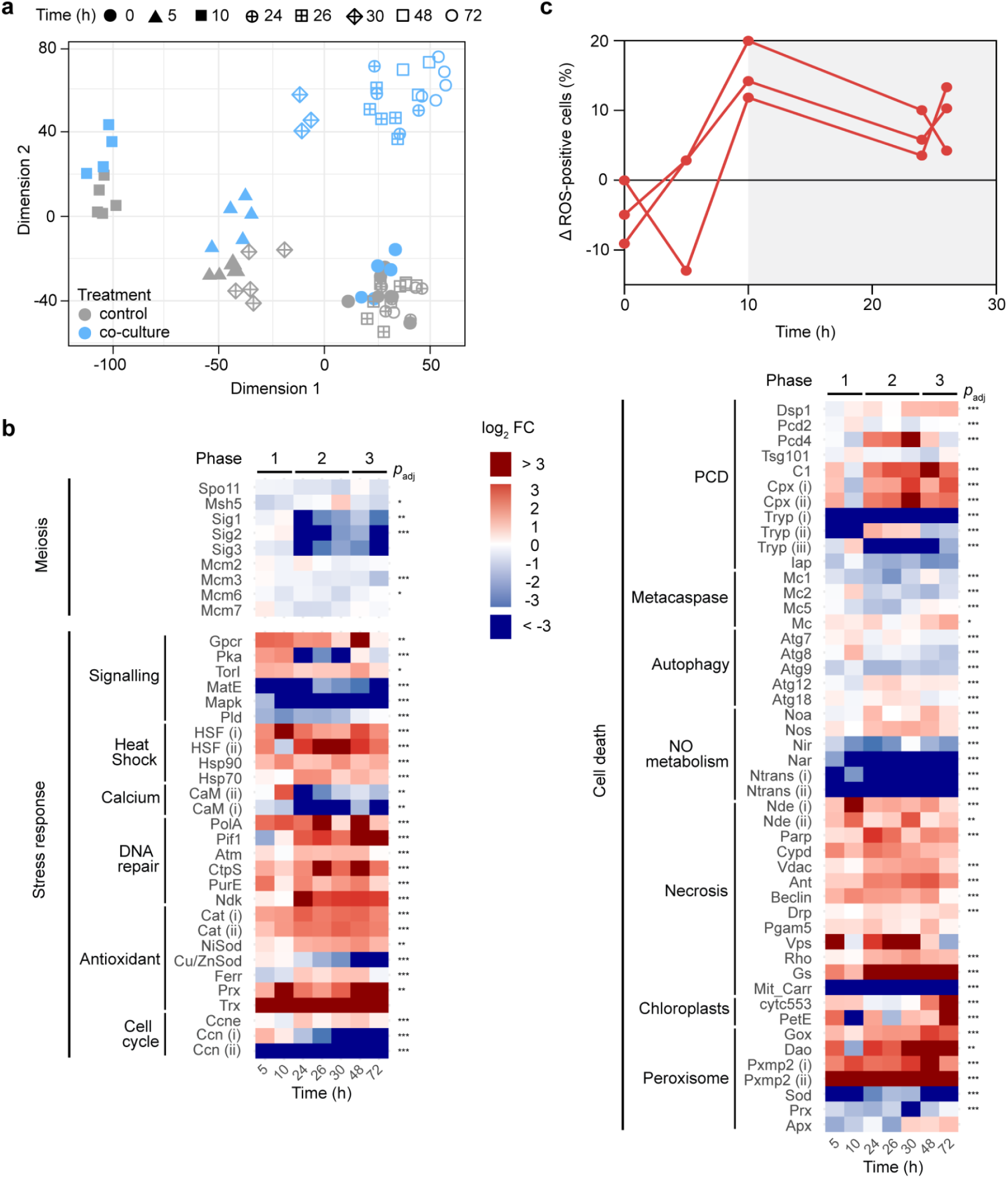
*T. rotula* in co-culture with *A. macleodii* displays transcriptomic signatures of cell death consistent with ROS-driven programmed necrosis. a Multidimensional scaling (MDS) plot based on variance-stabilising transformed (VST) RNA-seq data for *T. rotula* from 5 independent co-cultures with *A. macleodii* and 5 independent control cultures (treated with 1% 2216 MB). Each symbol represents one sample, with colours indicating control (grey) or co-culture (blue) and symbols indicating different time points (see legend). **b** Relative transcript abundances of *T. rotula* genes after 5, 10, 24, 26, 30, 48 and 72 h, given as log_2_-fold change (log_2_FC) relative to the 0 h time point and control cultures without *A. macleodii* (colour scale: ≤ –3 = dark blue, -3 to 0 = blue to white, 0 to 3 = white to red, ≥ 3 = dark red). Genes are grouped by cellular process based on KEGG and Gene Ontology (GO) annotations and manual curation from the literature. The significance of changes in expression over the entire time series is indicated with asterisks (DESeq2 likelihood ratio test, * = *p*_adj_ < 0.1, ** = *p*_adj_ < 0.05, *** = *p*_adj_ < 0.001). Benjamini–Hochberg false discovery rate-adjusted *p*-values for all genes are provided in Supplementary Table S3. **c** Difference in the proportion of ROS-positive *T. rotula* cells at 0, 5, 10, 24, 26 h in co-culture with *A. macleodii* in comparison to control cultures (no treatment), measured by CM-H₂DCFDA staining. Symbols show independent biological replicates, >10000 cells counted per replicate.

Given the morphological similarity between protoplasts and auxospores we examined transcription of markers of diatom sexual production^45–51^ (Supplementary Discussion), and found they were not differentially expressed or downregulated in co-culture (Fig. 3b), indicating sexual reproduction pathways do not underlie the observed morphology.

The transcriptomic results suggest that *T. rotula* rapidly initiates a stress response upon co-culture with *A. macleodii*, sequentially activating signaling and repair pathways. Major transcriptional changes occurred as early as 5 and 10 h (Phase 1) and persisted, including upregulation of signaling proteins such as G protein-coupled receptor, *Gpcr,* and kinases *Tor* and *Pka*, alongside strong downregulation of *Mapk*, inositol phosphate signaling components (*Pld*), sensor histidine kinases (*HK)* and *MatE* (Fig. 3b): repression of *MatE* may reduce signaling molecule export to reinforce stress signaling^52,53^. Heat shock factors (*HSF*), key stress regulators, and effector heat shock protein *Hsp90* were upregulated within 5 h, followed by *Hsp70* after 24 h, indicating a phased stress and proteostasis program. Calmodulin (*CaM)* genes were strongly downregulated from Phase 2, suggesting repression of Ca²⁺-dependent signaling. Genes for DNA damage repair and antioxidants to neutralize reactive oxygen species (ROS) were sequentially activated during Phases 1 and 2. Among the most strongly upregulated genes was a peroxiredoxin (*Prx*), which increased more than twentyfold by 10 h and over fiftyfold by 72 h, consistent with *Prx* induction to detoxify H₂O₂ reported in *Thalassiosira pseudonana*^54^. The strong downregulation of two Cyclin (*Ccn*) homologs, regulators of cell division and fate, from 5 h and 30 h, indicates cell cycle arrest. These transcriptional patterns reflect a rapid response to *A. macleodii* and sequential activation of stress-signaling, antioxidant defenses and DNA/protein repair pathways, promoting stress acclimation and survival.

From Phase 2 onwards, coinciding with the morphological change, *T. rotula*’s transcriptome showed signatures of phytoplankton programmed cell death (PCD)^29,54–57^, indicating that sustained stress overcame acclimation and shifted cell fate. Upregulated genes included dual stress/PCD regulators *Pcd4*, Ca²⁺-binding *Dsp1*, *Hsp90* and *Prx*, and a number of cysteine-endopeptidases that help to execute PCD (Fig. 3b): Among these was a cathepsin, *Cpx* (i), with 50% amino acid sequence identity (AA id) to the PCD-linked *P. tricornutum* Cpx^55^, and 38% AA id to human cathepsin Z, an executioner of the programmed necrosis pathway^58^. We also observed upregulation of *Cpx* (ii), showing 31% AA id to a human cathepsin L, and a *C1* peptidase, both of which encode an I_29 autoinhibitory domain common in regulated proteases associated with PCD. Moreover, expression of BIRC2-like inhibitor-of-apoptosis (*Iap*), which restrains cysteine endopeptidases, was significantly downregulated^54^. While several trypsin proteases linked to cell death in *T. pseudonana*^59^ were downregulated, this may indicate that protein catabolism becomes selectively regulated during the execution of cell death.

The activation of PCD markers did not fully align with canonical pathways. Key PCD genes showed weak or non-significant expression changes, including metacaspase enzymes (*Mc*)^60–64^, autophagy genes^65^, and genes whose expression is correlated with PCD in *S. marinoi*, *Pcd2* and *Tsg101*^57^ (Fig. 3b). Moreover, we found no clear evidence for nitric oxide (NO) signalling, a key regulator of PCD in other algae^66^ (Supplementary Discussion). These findings suggest *T. rotula* in co-culture with *A. macleodii* undergoes a metacaspase-, NO- and autophagy-independent form of PCD, thus divergent from canonical pathways studied in phytoplankton so far.

*T. rotula*’s PCD process displayed transcriptional signatures of mitochondrial ROS-linked PCD pathways, particularly programmed necrosis characterized in other eukaryotes but not previously described in algae^58,67,68^. Two alternative NADH dehydrogenases (Nde) were strongly upregulated at 10 h: *Nde* (i) (33% AA id to fungal *Nde-2*) increased by more than twentyfold at 10h, and carries an EF hand domain, consistent with calcium regulation (Fig. 3b). *Nde* (ii) (36% AA id to yeast *Nde1*; and to fungal *Nde-2*) also peaked at 10 h, coinciding with the commitment of *T. rotula* to protoplast-linked cell death. In budding yeast, Nde1 acts as a death effector^69^ and Nde1/2 in the fungus *N. crassa* directly generates mitochondrial ROS and promotes cell death^70^. Alternative NADH dehydrogenases are distant homologues of mammalian apoptosis-inducing factor (AIF), whose release from mitochondria triggers ROS-linked PCD pathways in mammals, including programmed necrosis^58,71,72^. Additional hallmarks of programmed necrosis in mammalian cells include PARP1 activation, which impairs respiration and promotes AIF release from mitochondria^73,74^, and the opening of the mitochondrial membrane permeability transition pore (mPTP), which further depolarises the mitochondria membrane, contributing to ATP depletion and ROS generation^67,75^. At 24 h *T. rotula* upregulated transcription of *Parp* and the three components of the mPTP; *Vdac, Ant* and *Cypd*. Putative Nde and mPTP involvement parallels mitochondrial ROS-linked cell death and programmed necrosis pathways described in other eukaryotes^58,68^. Mitochondrial stress is further suggested by induction of mitophagy regulators in *T. rotula* (*Beclin1*, *Drp, Pgam5, Rho, Vps)*^76^ and the sustained expression of glutamine synthetase (*Gs*), indicative of an attempt to mitigate redox imbalance. Transcriptomic signatures of mitochondrial collapse indicate that *T. rotula* is unable to sustain energy-dependent death pathways and undergoes a necrotic form of cell death.

ROS production appears to precede and potentially trigger the cell death process in *T. rotula*, with peroxisomes likely amplifying mitochondrial dysfunction. Strong regulation of thylakoid membrane alternative photosystem I (PSI) electron donors *PetE* and *Cyt-553* may indicate chloroplast redox imbalances (Fig. 3b). Upregulation of *Dsp1*, previously linked to cyclic electron flow around photosystem I (PSI) during cell death in *T. pseudonana*^54,56,77^, further supports altered chloroplast ROS metabolism. H_2_O_2_-producing peroxisomal enzymes D-amino acid oxidase (*Dao*) and glycolate oxidase (*Gox*), which plays a role in photorespiration and PCD in *Skeletonema marinoi*^57^, were upregulated from 5 h onward. Multiple *Pxmp*2 genes, encoding the H_2_O_2_-permeable peroxisomal membrane pore, were also strongly upregulated, with *Pxmp*2 (ii) expression remaining elevated more than eightfold at all time points. Concurrently, peroxisomal antioxidants such as *Sod*, *Prx* and ascorbate peroxidase (*Apx*) were not upregulated, suggesting elevated ROS production from peroxisomes and leakage into the cytosol. ROS staining with CM-H_2_DCFDA (Methods) showed that after 10 h of co-culture, the proportion of ROS-positive *T. rotula* cells was 8–15% higher than in the negative control, a significantly greater difference than at 0 h (Wilcoxon signed-rank test, *P* < 0.05; Fig. 3c), further supporting the involvement of ROS in the observed phenotype. Taken together, these results suggest that mitochondrial and peroxisomal-mediated oxidative stress drives a programmed necrotic-like cell death pathway in *T. rotula*, likely underlying the protoplast protrusion and loss of membrane integrity we have observed in co-culture with *A. macleodii*.

### Virulence strategies enabling nutrient extraction by *A. macleodii*

Of the 3740 genes transcribed by *A. macleodii* in co-culture, 2917 (78%) were differentially expressed in co-culture relative to the 0 h timepoint (likelihood ratio test, adjusted *P* < 0.1; Methods), with distinct profiles at each time point (Fig. 4a). Upregulation was observed in genes involved in motility, chemotaxis, and biofilm formation — virulence pathways enabling host colonisation^2,78,79^. In the KEGG flagellar assembly and chemotaxis pathways, 86% and 89% of genes were differentially expressed, with strong upregulation peaking in Phases 2 and 3, as illustrated by representative genes in Fig. 4b. These include Sigma-54 (*RpoN*), a regulator of virulence and flagella synthesis^80^, which was consistently upregulated in co-culture. 95% of biofilm- and attachment-related genes were differentially expressed, showing overall modest upregulation. Two of the strongest upregulated genes at 5 h were a BON-domain protein, upregulated more than thirtyfold that binds phospholipid membranes^81^ and a Big-9 domain protein upregulated more than sixfold, implicated in adhesion to hosts, including diatoms^82,83^ (Fig. 4b). The latter contains the COG2931 motif characteristic of Ca²⁺-binding RTX toxins, which show pore-forming, and cytotoxic activity^84^. In agreement with these findings, phase contrast and scanning electron microscopy (SEM) revealed bacterial biofilm formation and extracellular matrix on and around diatom cells (Fig. 4c).

**Figure 4.**
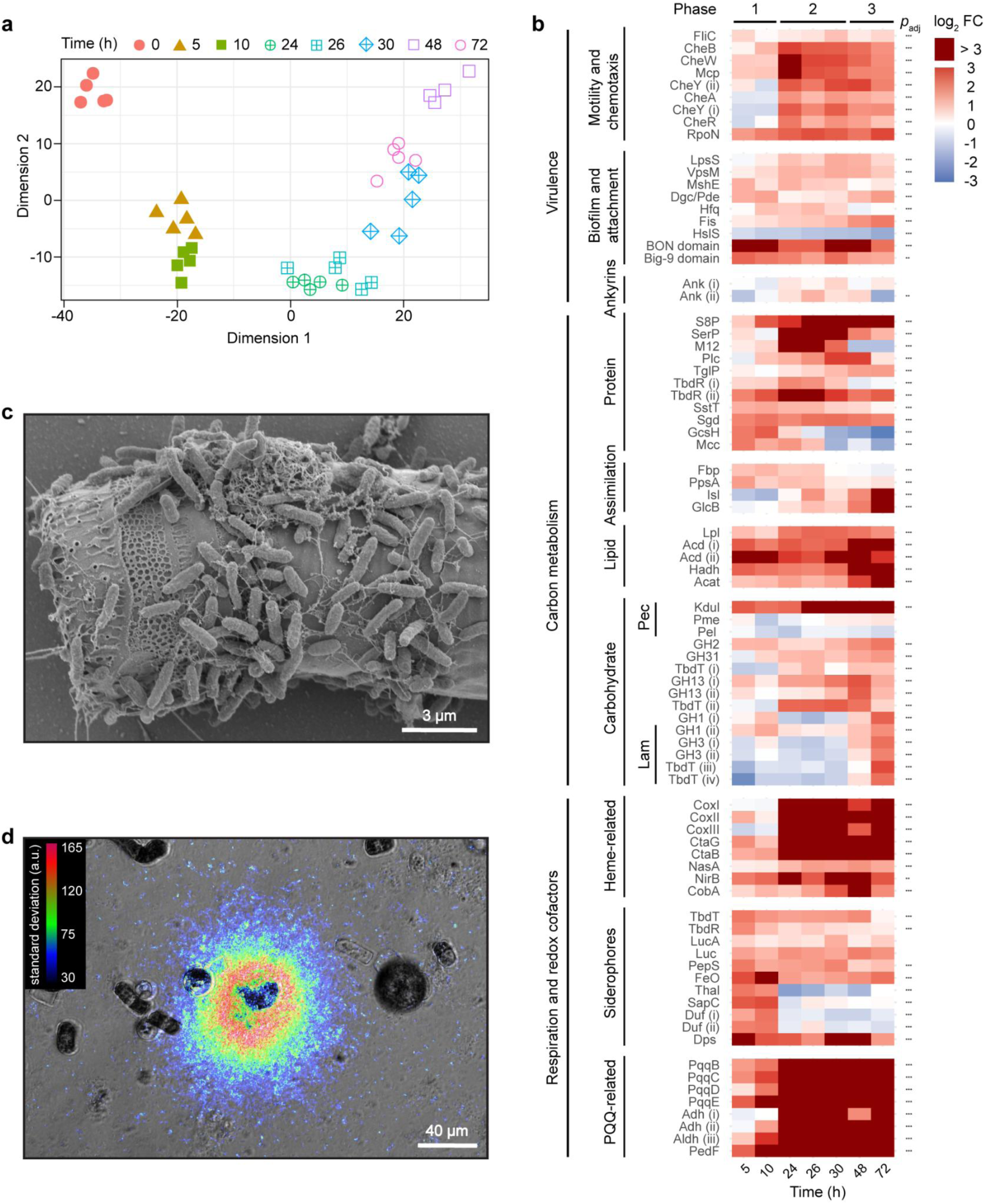
*A. macleodii*’s virulence strategy: coordination of motility, chemotaxis, biofilm formation and dynamic metabolism. **a** Multidimensional scaling (MDS) plot based on variance-stabilising transformed (VST) RNA-seq data for *A. macleodii* from 5 independent co-cultures with *T. rotula*; each symbol represents one sample, with colours and symbols indicating different time points (see legend). **b** Relative transcript abundances of *A. macleodii* genes after 5, 10, 24, 26, 30, 48 and 72 h in co-culture, gives as log_2_-fold change (log_2_FC) relative to the 0 h time point (colour scale: -3 to 0 = blue to white, 0 to 3 = white to red, ≥ 3 = dark red). Genes are grouped by cellular process based on KEGG annotations and manual curation from the literature. The significance of expression changes over the entire time series is indicated with (DESeq2 likelihood ratio test, * = *p*_adj_ < 0.1, ** = *p*_adj_ < 0.05, *** = *p*_adj_ < 0.001). Benjamini–Hochberg false discovery rate adjusted *p*-values for all genes are provided in Supplementary Table S5. **c** Scanning electron micrograph of a co-culture fixed after 24 h, showing biofilm formation by *A. macleodii* on the surface of a *T. rotula* cell. Scale bar 3 µm. **d** Time-lapse video microscopy image showing *A. macleodii* swarming around a lysing *T. rotula* cell (video available as Supplementary Movie 3). The image is a Z-projection of the standard deviation of pixel intensity across 45 frames (1.8 seconds), used as a proxy for bacterial motility, false-coloured and overlaid on a single video frame. Scale bar 40 µm.

**Figure 5.**
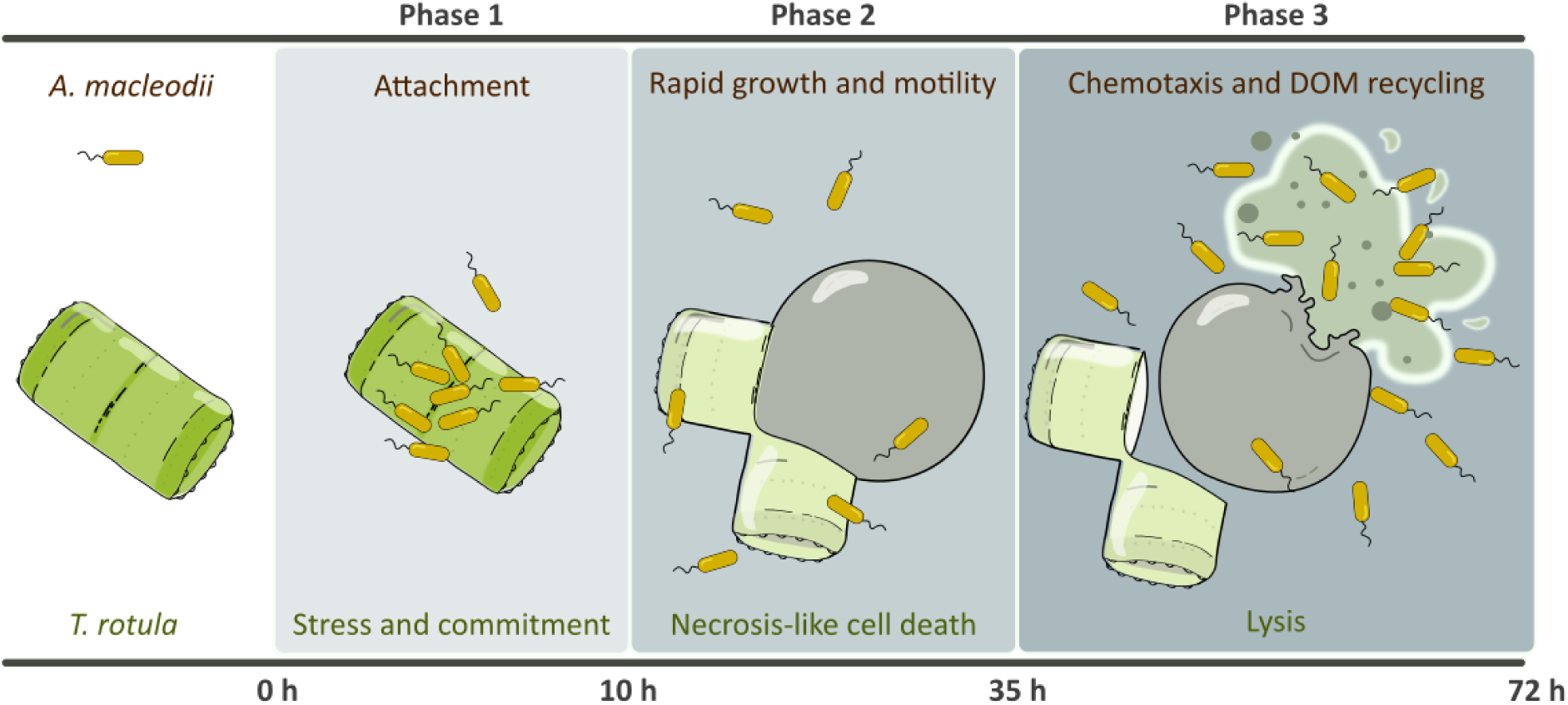
Conceptual model for the algicidal interaction between *T. rotula* and *A. macleodii*. The interaction unfolds in three distinct phases. In Phase 1 (0–10 h), *A. macleodii* attaches to *T. rotula*, triggering a transcriptional programme that includes stress-response genes and commits it to the terminal morphological change. Phase 2 (10-35 h) is marked by extrusion of the diatom’s protoplast from its silica frustule, and the upregulation of genes associated with necrotic cell death, including peroxisomal and mitochondrial pathways leading to reactive oxygen species (ROS) accumulation. Meanwhile, In Phase 3 (after 35 h), the exposed protoplasts, lacking the protective silica shell, become susceptible to lysis. *A. macleodii* then exhibits chemotactic swimming toward the extruded cellular material and upregulates genes involved in the degradation and assimilation of diatom-derived organic matter.

Gene expression data indicate that *A. macleodii*’s algicidal activity does not rely on canonical secretion pathways or common effectors, pointing to previously uncharacterized mechanisms of microbial interaction. Genes for the type II secretion system, Sec and Tat pathways, and major membrane vesicle proteins^85^ were not upregulated (Supplementary Table S6) and known eukaryotic-interference factors such as ankyrin repeat proteins^86^ exhibited minimal changes (Fig. 4b). Extracellular proteases (*S8P*, *SerP, M12, Plc*; Fig. 4b), shown to be effectors in other algicidal interactions^11,87,88^, were only induced after 24 h, well after *T. rotula* cells were committed to the morphological change and many had already undergone the transformation (Fig. 4b). Whilst chemical inhibition of proteases could not be tested due to toxicity to *T. rotula*, the gene expression data suggest that proteases are likely not the causative agents of protoplast formation, instead acting saprotrophically, via extracellular digestion, to support *A. macleodii*’s growth.

*A. macleodii*’s transcriptome reveals dynamic, phase-dependent utilization of algal derived compounds, enabling it to exploit the enriched substrate pool created by *T. rotula*’s breakdown. *A. macleodii* initially exploited proteinaceous carbon, with upregulation of extracellular proteases, peptide and amino acid transporters (*TbdR, SstT*), and catabolism enzymes (*Sgd*, *GcsH*, *Mcc*) (Fig. 4b). Modest induction of gluconeogenic enzymes (*Fbp* and *PpsA*) between 5-26 h, indicates protein-derived carbon supported anabolic growth during this period. As *T. rotula*’s membranes were exposed and degraded during Phases 2 and 3, *A. macleodii* upregulated lipid catabolism, upregulating lipases that liberate fatty acids (*Lpl*), β-oxidation (*Acd*, *Hadh*, *Acat)* and glyoxylate cycle enzymes (*Isl* and *GlcB)*, enabling the conservation of fatty-acid-derived carbon for biomass. *A. macleodii* utilized carbohydrates sparingly in co-culture overall (Supplementary Discussion; Supplementary Table S6) but upregulation of laminarin utilization genes in Phase 3 is consistent with release of this storage glucan during *T. rotula* lysis, and indicative of a broader nutrient ‘feast’ by 72 h (Fig. 4b). In parallel, *A. macleodii* leveraged redox cofactors to boost respiration of host-derived substrates, increasing investment in siderophores for iron acquisition (Supplementary Discussion), heme cofactor synthesis and heme-containing electron transport components. Furthermore, *A. macleodii* strongly induced synthesis of pyrroloquinoline quinone (PQQ), and PPQ-dependent alcohol dehydrogenases (*Adh*), coupling ATP production to oxidation of alcohols, including potentially methanol (Fig. 4b; Supplementary Discussion). Overall, *A. macleodii* dynamically adapted its metabolism to exploit the succession of diatom-derived substrates, culminating in a nutrient surge following *T. rotula* cell lysis.

## Discussion

### A novel algicidal strategy: exploiting protoplast formation

Our study uncovers a new strategy of bacterial exploitation of diatoms. The widespread heterotrophic bacterium *Alteromonas macleodii* exhibits algicidal activity against *Thalassiosira rotula* by inducing over 95% of cells in the population to emerge from their protective silica shells, exposing vulnerable protoplasts, which are then exploited by the bacteria. This interaction progresses through three distinct phases: *T. rotula* cells first undergo growth arrest (0-10 h), rapid and irreversible protoplast formation (10-35 h), and subsequent cell lysis (from 35 h). The high susceptibility of protoplasts to lysis, with 27% of the population lysed by 72 h, indicates that protoplast formation is a stress-induced, non-adaptive response, which *A. macleodii* exploits to fuel its own proliferation.

To the best of our knowledge, protoplast emergence in response to bacteria has not been previously reported in diatoms, establishing a novel algicidal mode. However, similar morphologies occur under nutrient limitation and stress. In stationary-phase *T. rotula* CCMP 3096 cultures (>10 days), 20% of cells exhibited protoplast formation in the absence of *A. macleodii* (Extended Data Fig. 1a). Protoplasts have also been noted in stationary-phase or stressed cultures of several other diatom species (M. Montresor, Naples; Personal communication), including *Coscinodiscus wailesii* CCMP 2513, *Thalassiosira pseudonana* CCMP 1335, *T. pseudonana* CCMP 3667, *Thalassiosira rotula* CCMP 3264, and *Thalassiosira weissflogii* CCMP 1051 (Extended Data Fig. 1b). These observations identify protoplast emergence as a general but poorly understood stress response, which *A. macleodii* may exploit to circumvent the diatom’s protective frustule. Stress accumulation at the end of algal blooms^29^ may heighten this vulnerability, underscoring the ecological relevance of this interaction.

To study this algicidal interaction and associated protoplast formation, we established co-culture conditions that reliably synchronized protoplast formation in *T. rotula* after 72 h. Diatoms were co-cultured with *A. macleodii* at 1 x 10⁶ cells/mL (approximately a 1:10 diatom-to-bacteria ratio), the upper range observed in nature. This also introduced 1% (v/v) 2216 MB medium into the co-culture which, based on previous reports of DOM-activation^11^, may have stimulated the algicidal activity of *A. macleodii*. Indeed, addition of 1% 2216 MB alone to xenic *T. rotula* cultures significantly increased protoplast formation (Fig. 1c), though to a much lesser extent than *A. macleodii* co-addition. As *A. macleodii* is a known member of *T. rotula*’s microbiome^44^ and was detected in our xenic diatom cultures, this effect likely reflects stimulation of resident *A. macleodii* to induce the phenotype, rather than a direct diatom response. These findings highlight an environmentally relevant scenario in which DOM-activated *A. macleodii* during bloom conditions could trigger protoplast formation in diatoms.

### Evidence for programmed necrotic-like PCD in *T. rotula*

Our results demonstrate that the death of *T. rotula* cells during co-culture with *A. macleodii* is a programmed response to prolonged stress, rather than a passive consequence of bacterial attack. This marks the first evidence of PCD in *T. rotula* and underscores its role in bacteria-phytoplankton interactions^89^. Commitment to protoplast emergence within 9 h, well before the morphological transformation begins, together with upregulation of a subset of PCD markers^29,54,57^, supports activation of a distinct PCD pathway. While metacaspase homologues^60–64^ were not induced, *T. rotula* strongly upregulated a cathepsin Cpx and related proteases, whose homologues in *Phaeodactylum tricornutum* and *Chlamydomonas reinhardtii* mediate oxidative stress-induced PCD^55^. This remodelling of proteolytic transcription indicates activation of a redox-sensitive, metacaspase-independent pathway, thus distinct from canonical phytoplankton PCD pathways.

The programmed death of *T. rotula* exhibits phenotypic hallmarks of necrosis - perinuclear organelle clustering and plasma membrane rupture - together with genotypic signatures resembling programmed necrosis pathways described in other eukaryotes. The strong transcriptional spike of two alternative NADH dehydrogenase genes (Nde) at 10 h coincided with elevated ROS detected in *T. rotula* in co-culture, consistent with their role as mitochondrial ROS-generating death effectors in fungi^69,70^. As *Nde* is a distant homologue of the mammalian apoptosis-inducing factor (AIF), they have been proposed to serve as redox-regulated cell death switches, conserved across eukaryotes^68–70^. Therefore, the upregulation of *Nde* in *T. rotula* may suggest mechanistic similarities with AIF-mediated programmed necrosis studied in mammals^58,72^. In line with this interpretation, *T. rotula* exhibited additional hallmarks of this pathway^67,73,90^, including upregulation of *Parp,* components of the mPTP, markers of mitochondrial dysfunction, and evidence of both ROS and calcium involvement. In addition to mitochondrial ROS production, peroxisomal H₂O₂ production and chloroplast redox perturbations may contribute to oxidative stress amplification in this form of PCD, consistent with organelle-specific oxidative stress patterns regulating PCD in phytoplankton^91^. Calcium signaling, previously implicated in distinct phytoplankton PCD pathways^92^, is supported here by the induction of calcium-regulated genes in Phase 1 (e.g. *Nde*(i)) and downregulation of calmodulins in Phase 2, which may reflect an adaptation to prolonged calcium signaling. Overall, we present evidence for an alternative PCD pathway in *T. rotula*, characterized by necrotic phenotypes and molecular parallels to programmed necrosis in other eukaryotes. Protoplast formation observed here, and reported under stress in other diatom species, may represent a conserved morphological feature of this pathway. These findings add to evidence for non-apoptotic death programs in algae^93,94^ and more broadly support the existence of a conserved programmed necrosis-like pathway with deep evolutionary origins in eukaryotes^68^.

### Mechanistic insights into *A macleodii*’s algicidal lifestyle

Protoplast induction was unique to *A. macleodii* and its close relative *A. mediterranea*, and the effect was restricted to *T. rotula* among the species tested, with the strain-specific magnitude suggesting a targeted algicidal mechanism and potentially a co-evolutionary relationship. Such specificity has important ecological implications, as targeted antagonism can govern *T. rotula* bloom decline^33^ and restructure microplankton community composition. In contrast to the algicidal interaction described here and elsewhere^11^, *A. macleodii* is not exclusively antagonistic but exhibits remarkable divergence across the species complex, ranging from opportunistic copiotrophy^35,38,40^ to beneficial associations that promote phytoplankton growth^95–98^. In the interaction studied here, *A. macleodii* repurposed its copiotrophic toolkit, upregulating motility, chemotaxis, biofilm formation, attachment and metabolic flexibility, to mediate host exploitation in a coordinated attack-and-consume strategy. The ability to shift its ecological function from opportunistic copiotroph to predatory bacterium opens up new niches for *A. macleodii*: rather than passively scavenging the decay of a bloom, *A. macleodii* can be recategorized as an algicidal bacterium that actively generates sources of energy and carbon through host manipulation, potentially contributing to bloom termination.

Investigating the basis of *A. macleodii’s* antagonistic activity, we found no evidence for conventional mechanisms, suggesting novel, uncharacterized algicidal effectors are at play. The mechanism is contact-dependent and strong candidates include attachment genes, particularly the upregulated RTX-like toxin, implicated in adhesion to diatom hosts and cytotoxic activity^82,83^. The induction of PCD in *T. rotula* may reflect a multifactorial strategy whereby the bacterium manipulates latent host stress pathways, possibly explaining the difficulty to pinpoint a single effector. Future work should aim to resolve whether protoplast formation represents an involuntary stress response or a finely tuned bacterial manipulation, as well as the molecular underpinnings of this highly effective algicidal mechanism.

### Ecological implications

The presence of a programmed necrotic-like cell death pathway, involving protoplast emergence from the frustule and heightened vulnerability to lysis, has broad implications for our understanding of *T. rotula*’s life history and population dynamics. The ability of *A. macleodii* to induce this algicidal response highlights the seismic impact bacterial interactions can exert on diatom ecology. This interaction has cascading effects on ocean biogeochemistry: by impairing productivity, promoting diatom lysis and removing the ballast provided by the silica frustule, it likely reduces the export of carbon and nutrients to deeper waters. *A. macleodii* has also been shown to form biofilm aggregates that reduce particle sinking and further alter carbon export efficiency^99^; this bacterium can thus reshape microbial communities and nutrient cycles, paralleling the well-documented role of viruses^100–102^. Our findings underscore the far-reaching ecological consequences of a distinctive algicidal adaptation in bacteria, highlighting the tight coupling between microbial interactions and the oceanic carbon cycle.

## Methods

### Diatom culture

*T. rotula* CCMP 3096 cells were obtained from NCMA (Bigelow Laboratory) as xenic cultures, with a natural complement of bacteria. The cells were maintained in L1 enriched seawater medium (35 gL−1 instant ocean sea salt), at 14 °C with 12:12 day:night cycle and diluted at least every 14 days, or more frequently as required. *T. rotula* CCMP 3096 was used for all experiments except where explicitly stated to involve strains CCMP 3362 or CCMP 3264, obtained from NCMA and maintained following the same protocol (except maintaining cells at 22 °C for *T. rotula* CCMP 3362). *Phaeodactylum tricornutum* CCMP 2560, *Thalassiosira pseudonana* CCMP 3367 and *Thalassiosira weissflogii* CCMP 1050 were cultured following the same protocol as *T. rotula* (except maintaining cells at 18 °C for *P. tricornutum)*.

### Cell counting

Culture density was measured using a flow cytometer (cytoFLEX, Beckman Coulter) with all gain and threshold settings on automatic. Gating was performed based on diatom cell autofluorescence in three different channels of excitation, APC (638 nm), PE (585 nm) and FITC (525 nm), plotted over forward scatter (FSC). The background signal from L1 medium within these gates was confirmed as zero, and the average cell count across the three channels was taken as the number of cells per mL.

### Axenic cultures

Axenic *T. rotula* CCMP 3096 cultures were generated with the following protocol, which was verified to have no detrimental effect on diatom growth. A 5 mL sample of diatom culture in the exponential growth stage was filtered onto a 3 μm pore-size cell strainer and washed with 5 mL sterile L1 medium. 5 mL of L1 medium containing 20 µg/mL Triton-X 100 detergent (Sigma-Aldrich) was added to the cells for 1 min, then the liquid was removed and the cells washed with 5 mL medium. Diatom cells were then aspirated off the membrane and diluted in 10 mL medium containing the antibiotics gentamycin sulfate 67 μg mL⁻¹, ciprofloxacin 20 μg mL⁻¹, chloramphenicol μg mL⁻¹, ampicillin 10 μg mL⁻¹ and streptomycin sulfate 50 μg mL⁻¹ (all Sigma-Aldrich). Cells were incubated in this antibiotic-containing medium for 48 h under regular growth conditions and then transferred 1:10 to antibiotic-free medium. Bacterial decontamination was verified by transferring 100 μL of the diatom culture to 2 mL 2216 MB (BD Difco) and confirming that there was no change in the OD600 after 24 h (CO8000 Cell Density Meter, LabGene scientific).

### Bacterial culture

A stock of *A. macleodii* ATCC 27126 was obtained from the ATCC culture collection and stored at -80 °C as a glycerol stock. For experiments, 5 mL 2216 MB was inoculated from the glycerol stock and grown at 27 °C, 200 rpm for 12 h. Before addition to diatom cultures, the OD600 was measured using a spectrophotometer blanked with 2216 MB (CO8000 Cell Density Meter, LabGene scientific).

### Diatom and bacteria co-culture

The conditions for addition of *A. macleodii* to *T. rotula* were optimised and these standards followed in all experiments, except where explicitly stated to differ for experimental purposes. *T. rotula* cultures split in L1 medium 3 days prior and with concentrations between 1×10^4^ and 20×10^4^ cells mL^−1^ were used to ensure exponential growth and optimal imaging. Exponentially growing *A. macleodii* cells, seeded 12 h prior, were added to a final ratio of 1% (v/v) directly from the culture in 2216 MB. Each experimental and control condition was conducted in triplicate and the cultures were maintained in 24-well glass bottomed plates (Cellvis), allowing microscopic analyses to be performed directly. The plates were incubated at 14 °C with 12:12 day:night cycle, and for end point experiments images were taken at 0 h and 72 h (10X Objective, Nikon ECLIPSE Ti Microscope equipped with Hamamatsu ORCAFlash4.0 digital camera). The most accurate method for analysing protoplast morphologies proved to be manual, thus *T. rotula* cell types were categorised and counted by eye from the images. For time course experiments quantifying temporal dynamics and diatom cell lysis, plates remained on the microscope stage, with an LED panel constructed to provide a 12:12 day:night cycle (P3030W, LED express). Images were taken every 20 min for 72 h (10X Objective, Nikon ECLIPSE Ti Microscope equipped with Hamamatsu ORCAFlash4.0 digital camera). The resulting videos were analysed by eye, categorising the cells in each frame as cylindrical, spherical or lysed.

### Incubation experiments

To culture cells in a shared medium but in the absence of physical contact, sterivex filter units with 0.45 μm pore membranes were used (Merck, SVHVL10RC); *T. rotula* only cultures were injected into one side of the filter unit, and *A. macleodii*-*T. rotula* co-cultures into the other, which were then maintained at 14 °C with 12:12 day:night cycle. Positive control filter units were set up with co-cultures in both compartments and a negative control with *T. rotula* only cultures in both compartments. After 72 h, an aliquot from each compartment was placed on a glass slide and viewed under the microscope to quantify the cell phenotypes (10X Objective, Nikon ECLIPSE Ti Microscope equipped with Hamamatsu ORCAFlash4.0 digital camera).

### Scanning electron microscopy

For scanning electron microscopy (SEM), co-cultures were set up between axenic *T. rotula* and *A. macleodii* following the standard protocol and samples were fixed with glutaraldehyde at a final concentration of 2.5% at 0, 5, 24 and 48 h after addition of the bacteria. Fixed cells were attached to a silicon support coated with poly-L-lysine. The samples were then washed sequentially for 5 min with 5 mL 2.5% glutaraldehyde, 5 mL seawater, 5 mL 1% osmium tetroxide, and then again with 5 mL seawater. The samples were dehydrated with an ethanol series in distilled water, with 5 min at each concentration (30%, 50%, 70%, 90% and 100%) followed by three 5 min washes with water-free 100% ethanol. The material was critical point dried (Autosamdri-931, Tousimis) and the supports were mounted on aluminium stubs using silver paint. The sample was then uniformly metalized with 4 nm Pt-Pd by cathode sputter-coating (CCU-010 HV, Safematic). Cells were imaged using an SU5000 scanning electron microscope (HITACHI).

### Antibiotics and inhibitors

To determine the time point at which *T. rotula* becomes committed to the morphological change during co-culture with *A. macleodii*, antibiotics were added at incremental delays to inhibit *A. macleodii*. Final concentrations, confirmed to inhibit *A. macleodii* growth, were 50 μg mL⁻¹ kanamycin sulfate and 25 μg mL⁻¹ chloramphenicol; 100 μg mL⁻¹ ampicillin, which is non-inhibitory to *A. macleodii*, was included as a control (all Sigma-Aldrich). Antibiotics were added 0, 3, 6, 9, 24, or 48 h after co-culture initiation under standard conditions and the proportion of *T. rotula* protoplasts was quantified after 72 h.

### Dual Transcriptomics

#### Co-culture and RNA extraction

For total RNA extraction, *T. rotula* cultures in exponential growth phase (10 replicates; 560 mL at 1 × 10⁴ cells mL⁻¹ in 2 L Erlenmeyer flasks) were maintained at 14 °C under a 12:12 h light:dark cycle. Five treatment cultures were inoculated with 5.6 mL *A. macleodii* (12 h culture in 2216 MB; final 1% v/v), and five controls received an equal volume of sterile 2216 MB. Samples were taken from each of the 10 cultures at 0, 5, 10, 24, 26, 30, 48, and 72 h post-inoculation. At each time point, 500 μL was transferred to a 24-well glass-bottom plate for quantifying diatom phenotypes, and two 35 mL aliquots were harvested for RNA extraction by centrifugation (2,000 g, 3 min; Eppendorf 5804R, A-4-44 rotor). The pellets were resuspended in 500 μL ROTI Aqua-Phenol (Carl Roth), transferred to tubes containing glass beads (BashingBeads, Zymo Research), shaken (3 min, vortex setting 10; Vortex-Genie 2 with Multi Tube Holder, LabGene Scientific), incubated at room temperature for 2 min, then centrifuged at 12,000 g for 7 min (Eppendorf 5425R, FA-24 × 2 rotor). Supernatants were stored at −80 °C until processing of all time points.

#### Clean up, DNase, RT and qPCR

The RNA samples were cleaned and concentrated using an RNeasy Minikit (QIAGEN), following the RNA clean-up protocol, and eluting in 46 μL RNase-free water (kit-supplied). RNA concentration and purity were measured on a NanoDrop spectrophotometer (NanoDrop OneC, Thermo Fisher Scientific). All samples with an A260/A230 ratio below 1.80, indicating phenol contamination, were subjected to standard ethanol precipitation: 0.1 volumes 3 M sodium acetate (pH 5.2) and 3 volumes ice-cold 100% ethanol were added, samples were vortexed, and RNA was precipitated overnight at −80 °C. Thawed samples were centrifuged at 15,900 g for 30 min at 4 °C (Eppendorf 5425R centrifuge, FA-24 × 2 rotor), the pellets were washed twice with 0.5 mL ice-cold 75% ethanol (centrifuging at 4 °C for 10 min each time), air-dried, and then resuspended in 46 μL nuclease-free water. RNA concentration and purity were measured again on a NanoDrop OneC.

DNA digestion was performed with TURBO DNase (Thermo Fisher Scientific) in 50 μL reactions containing 5 μL 10X reaction buffer, 1 μL TURBO DNase, and 44 μL RNA solution, incubated at 37 °C for 30 min. Digestion was terminated with 5 μL DNase inactivation reagent, incubated at room temperature for 5 min, and centrifuged at 10,000 × g for 1.5 min. The supernatant was transferred to a fresh tube, and RNA concentration was measured again using both NanoDrop and the Qubit Broad-Range RNA Assay Kit (Thermo Fisher Scientific) in a 384-well plate format. In order to confirm gDNA removal and the presence of RNA, reverse transcription was performed in 20 μL reactions containing 15 μL nuclease-free water, 1 μL of 1:10 diluted RNA, and 4 μL 5X Reaction Mix (iScript RT supermix, Bio-Rad). For each sample, both RT and no-RT controls were prepared. Amplification of *T. rotula* and *A. macleodii* DNA was detected using the 7500 FAST Real-Time PCR System (Thermo Fisher Scientific) in reactions containing 4 μL 5X HOT FIREPol EvaGreen qPCR Mix Plus (Solis BioDyne), 250 nM each of *T. rotula* Rpl13 primers (Rpl13-F ACTTCGCCTCAAGCCAAAC, Rpl13-R TTGACCTCAGCCAATTCTCC) or *A. macleodii* 16S primers (16S-27F AGAGTTTGATCMTGGCTCAG, 16S-1492R GGTTACCTTGTTACGACTT), and 1 μL template solution. gDNA removal was verified by checking for the absence of amplification in no-RT reactions; RNA presence was confirmed by checking for amplification in RT reactions. Negative template controls were included for each primer pair. All reactions were conducted in duplicate. The PCR amplification program consisted of an initial denaturation step at 95 °C for 12 min, followed by 40 cycles of 95 °C for 15 s, 62 °C for 30 s, and 72 °C for 60 s. Melt curves were checked for each sample to confirm amplification of the correct product.

#### rRNA depletion and library preparation

rRNA depletion and dual-library preparation were performed using the Zymo-Seq RiboFree Total RNA Library Kit (R3003, Zymo Research), a probe-free method that generates combined libraries from both organisms. For each sample, 500 ng RNA from the post-DNase step was used as input. Dual-library concentration was measured with the Qubit dsDNA HS Assay Kit (Thermo Fisher Scientific), and size distribution and quality were assessed using the High Sensitivity D1000 ScreenTape assay on a TapeStation system (Agilent). An equimolar pool of all libraries was prepared and re-analysed using Qubit and TapeStation assays.

#### Sequencing

The dual RNA-seq libraries were sequenced as 150 bp paired-end reads using the NovaSeq Reagent Kit v1.0 (300 cycles) on a NovaSeq 6000 System (Illumina) at the Genetic Diversity Centre, ETH Zurich. Preliminary experiments confirmed that the ratio of reads mapping to *T. rotula* and *A. macleodii* was favorable, providing sufficient coverage of both organisms without requiring deeper sequencing.

### RNA sequence processing

The scripts used for transcriptome processing in this study are available on the GitHub page https://github.com/ClaraMarPe/T.rot_A.mac_transcriptome.

Sequence reads were processed to remove Illumina-specific adaptors and were quality-trimmed using BBduk (BBtools v37.61; sourceforge.net/projects/bbmap) with default parameters and a minimum length of 100 nucleotides. Error correction of the reads was performed using the Bayes-Hammer module of the SPAdes assembler^103^ (v3.15.4). Only paired reads were retained, and only those pairs where both reads mapped in the correct orientation and with the correct insert size. To remove ribosomal RNA-like sequences from the dataset, high-quality reads were mapped sequentially to the SILVA SSU and LSU databases^104^(v138.1) and to the Y2K 5S rRNA database^105^, using BBmap v37.36 (sourceforge.net/projects/bbmap). Reads with a sequence identity threshold of >70% were removed.

To obtain an overview of the microbial community composition in the xenic *T. rotula* cultures (without co-cultivation with *A. macleodii*), reads that had been mapped to small subunit (SSU) rRNA during the rRNA-filtering step were mapped to the SILVA non-redundant SSU Ref database (v.138.1) and assigned to an approximate taxonomic affiliation (nearest taxonomic unit, NTU) using PhyloFlash v3.0^106^ (http://github.com/HRGV/phyloFlash). The results provided a qualitative estimate of the community composition, as residual rRNA reads were obtained from libraries subjected to rRNA depletion. *Alteromonas macleodii*–assigned NTUs were detected (representing ∼3.1% of the residual rRNA reads across four replicates), confirming its presence in the xenic *T. rotula* CCMP 3096 cultures. Results are summarized in Supplementary Table S1.

### *T. rotula* de novo transcriptome assembly and annotation

Due to the lack of a reference genome for *T. rotula* at the time of analysis, *de novo* assembly of the diatom transcriptome from the dual RNA-seq data was performed. To facilitate this, mRNA reads resembling *A. macleodii* were removed by mapping to the *A. macleodii* ATCC 27126 genome (NCBI:txid529120) at low stringency (>70% sequence identity) using BBMap. The remaining reads were normalized *in silico* to a target average depth of 80× using BBnorm (BBTools v37.61). A *de novo* transcriptome co-assembly was generated from the normalized, concatenated reads of all samples using rnaSPAdes^107^ (SPAdes v3.15.4) with default parameters (K-mer sizes: 49, 73). Non-diatom contaminants were removed by DIAMOND BLASTx (v2.0.13) searches against the NCBI non-redundant protein database (nr; July 2022), discarding sequences with significant hits (e-value <= 10^-15) to Bacteria, Archaea, or Metazoa. To reduce transcriptome redundancy, assembled transcripts were clustered using CD-HIT-EST^108^ (v4.8.1) with a sequence identity threshold of 95% and global alignment (-c 0.95 -g 1). The decontaminated transcriptome assembly comprised 76,185 contigs (total length 85.4 Mb; N50 = 2,231 bp; average contig length = 1,122 bp). CDS retained after filtering had a mean length of 1.3 kb and N50 of 1.8 kb (n = 35,514).

Quality and completeness of the *de novo* transcriptome assembly were evaluated using Benchmarking Universal Single-Copy Orthologs (BUSCO)^109^ v5.8.3 with the Stramenopiles lineage dataset, yielding 97% complete BUSCOs (59% single-copy, 38% duplicated), 2% fragmented, and 1% missing out of 100 orthologs. To further evaluate concordance with genomic resources, transcript-derived coding DNA sequences (CDS; see below) were aligned to the recently published *T. rotula* genome (strain FE80)^110^, and reciprocal best hits were identified. At permissive thresholds (≥50 % identity and ≥50 % coverage), 95 % of transcriptome-derived CDS and 80 % of genome CDS showed a best hit to the other assembly, while under more stringent criteria (≥90 % identity, ≥90 % coverage), 66 % of transcript-derived CDS still aligned to genome CDS. Reciprocal best-hit analysis identified ∼47 % of transcripts as having a one-to-one orthologous counterpart in the genome, and ∼32-43 % when identity and coverage filters were applied. A summary of assembly statistics, BUSCO scores, and genome alignment metrics is provided in Supplementary Table S2.

Transdecoder (https://github.com/TransDecoder/TransDecoder/) v.5.7.1 was used to predict candidate coding regions (CDSs), retaining only the single best open reading frame (ORF) per transcript. To reduce false-positive ORF predictions, we applied the --retain_blastp_hits and --retain_pfam_hits options during TransDecoder.Predict, using homology searches against the NCBI nr protein database (DIAMOND BLASTp, July 2022) and conserved domain searches against Pfam (HMMER hmmsearch, v3.3.2). Transcript sequences were translated into protein sequences, using a minimum ORF of 100 amino acids (TransDecoder.LongOrfs). Functional annotation of the translated protein sequences was performed with DIAMOND BLASTp against the NCBI and UniProt nr databases (both accessed July2022), and domain prediction using InterProScan (v5.53-87.0)^111^ against the InterPro families database (v89.0)^112^. Additionally, orthology prediction was performed with eggNOG-mapper (v2.1.6)^113^ against the EggNOG database (v5.0).

### Quantification of transcript expression and differential expression analysis

Quantification of gene expression was performed by mapping the (non-normalized) mRNA reads to either the *de novo* assembled contigs of *T. rotula* or to the reference genome of *A. macleodii* (GenBank: CP003841.1) using Bbmap with mapping identity threshold of 97% (minid=0.97). Alignments below this threshold were filtered out (idfilter=0.97).Mapped reads were assigned to predicted ORFs in *T. rotula* and to the existing, annotated coding sequences of the reference *A. macleodii* genome using featureCounts^114^ (Subread package, v2.0.3), generating gene-level count tables that were used for downstream processing in R v 4.3.3. On average, 60.85 % (range 27.29-87.34 %) of all mRNA reads mapped to the *T. rotula* CDS dataset across all samples. In co-culture samples, an average of 8.62 % (range 2.85-37.7 %) of reads mapped to the *A. macleodii* CDS. Transcript representation and coverage for both the *de novo* transcriptome assembly and the reference *A. macleodii* genome are summarized in Supplementary Table S2.

Differential expression analyses were performed separately on the mapped read counts for *A. macleodii* and *T. rotula* using DeSeq2^115^(v1.42.1). Only genes with a read count of 10 or higher in at least three samples were included in the analysis. Hierarchical clustering of the samples was performed using DESeq2-normalized data (subjected to Variance Stabilizing Transformation, VST) using Euclidean distance as the similarity metric. Differential expression of *T. rotula* genes between control (xenic *T. rotula* cultures) and *T. rotula* with *A. macleodii* co-culture samples was assessed at each time point (5 h, 10 h, 24 h, 26 h, 30 h, 48 h, 72 h) using a likelihood ratio test in DESeq2. A negative binomial generalized linear model (GLM) was fitted using the design formula “∼treatment+time+treatment:time”, an approach that controls for variation among replicates at time 0 h and tests whether the temporal expression patterns differ between treatments. Differential expression in *A. macleodii* upon interaction was assessed exclusively in co-culture samples. Pairwise comparisons were made between each timepoint and time 0 h, using a simpler design formula, “∼time” in DESeq2. For all analyses, statistical significance was determined using a Benjamini–Hochberg false discovery rate (FDR) threshold of 5%.

## Acknowledgments

We thank R. Naisbit for help with editing the manuscript and Oliver Müller and Johannes Keegstra for valuable discussions. Computational analyses were performed using the Euler high-performance computing cluster at ETH Zürich. Data produced and analysed in this paper were generated in collaboration with the Genetic Diversity Centre (GDC), ETH Zurich. We acknowledge the Scientific Center for Optical and Electron Microscopy (ScopeM), ETH Zürich, for expert support in electron microscopy.

## Author contributions

I.S, C.M.P, R.S and U.A designed research. I.S, C.M.P, and U.A conducted microbiology and microscopy experiments. I.S, C.M.P, and U.A analyzed and interpreted data. I.S, C.M.P, R.S and U.A wrote the paper.

## Data availability

The quality-filtered mRNA reads and the *Thalassiosira rotula* de novo transcriptome assembly used as a reference for transcript mapping are available in the European Nucleotide Archive under the project number PRJEB72406. The *Alteromonas macleodii* ATCC 27126 complete genome used for the bacterial transcript mapping is publicly available under GenBank accession number CP003841.1.

## Extended Data

**Extended Data Figure 1.**
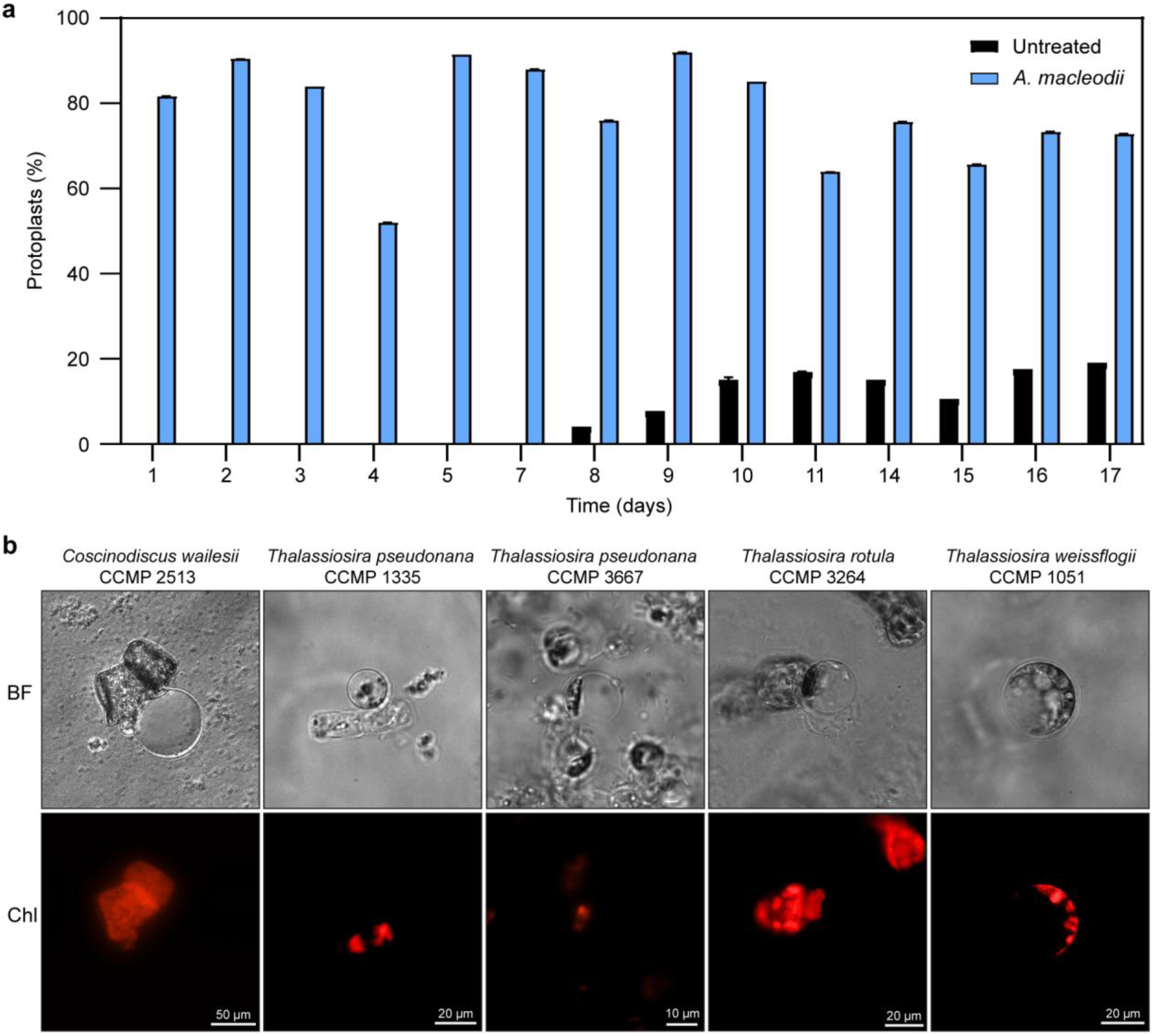
Naturally occurring protoplast formation in *T. rotula* and other diatoms. **a** Proportion of *T. rotula* CCMP 3096 cells forming protoplasts. A diatom culture was maintained for 17 days, and on each day indicated, aliquots were taken and incubated for 72 h either untreated (black) or with *A. macleodii* (blue); >100 diatom cells were scored by phase-contrast microscopy after 72 h incubation. **b** Light microscopy (BF) and corresponding chlorophyll autofluorescence (Chl) images showing protoplasts in five diatom species: *Coscinodiscus wailesii* CCMP 2513, *Thalassiosira pseudonana* CCMP 1335, *T. pseudonana* CCMP 3667, *Thalassiosira rotula* CCMP 3264, and *Thalassiosira weissflogii* CCMP 1051.

