## Supplementary Information for "Breaching the diatom frustule: *Alteromonas macleodii* triggers protoplast emergence and programmed necrotic-like cell death in the oceanic diatom *Thalassiosira rotula*"

Affiliations:
[1] Institute of Environmental Engineering, Department of Civil, Environmental and Geomatic Engineering, Eidgenössische Technische Hochschule Zürich (ETHZ), Stefano-Franscini-Platz 5, 8093 Zürich, Switzerland.

[2] Current address: Limnological Research Station, Department of Hydrology; University of Bayreuth, Universitätsstraße 30, Bayreuth, Germany.

[3] Department of Plant and Environmental Sciences, Institute of Life Sciences, The Hebrew University of Jerusalem, Jerusalem, Israel.

**This PDF file includes:**

**Description of Supplementary Tables S1–S6**

**Description of Supplementary Videos S1–S3**

**Supplementary Text**

### Description of Supplementary Tables

**Supplementary Table S1. (Excel Workbook)**
Taxonomic composition of SSU rRNA reads recovered from xenic *T. rotula* CCMP 3096 cultures (replicates S9-S12, representing the control condition in the metatranscriptome analysis) at time 0 h. Reads were classified with PhyloFlash v3.0 against the SILVA SSU Ref NR99 database (v138.1). Values represent read abundance and relative proportion (%) of each taxonomic lineage. Due to prior rRNA depletion, results provide an approximate (non-quantitative) indication of the native bacterial community present in the diatom cultures.

**Supplementary Table S2. (Excel Workbook)**

Quality assessment of de novo *T. rotula* CCMP 3096 transcriptome assembly, containing the following tabs:

**Assembly stats:** Separate summaries are shown for the complete transcriptome assembly and for the subset of predicted CDS regions used in downstream analyses. The metrics include total sequence length (sum, in base pairs), number of sequences (n), average and maximum contig lengths (in bp), and standard N-statistics (N50–N100). Ncount shows the total number of ambiguous nucleotides (N's) in the assembly, and Gaps indicates the number of gap regions detected.

**BUSCO:** Benchmarking Universal Single-Copy Orthologs completeness assessment of the *T. rotula* de novo transcriptome assembly and predicted CDS dataset, evaluated against the Stramenopiles_odb10 lineage dataset (BUSCO v5.8.3 in *transcriptome* mode). Report includes the counts of Complete (C), Single-copy (S), Duplicated (D), Fragmented (F), and Missing (M) BUSCOs out of the total number of n ortholog groups searched.

**RBHs:** Summary of reciprocal best-hit (RBH) analyses across varying sequence identity (50–90%) and alignment coverage (50–90%) thresholds between *T. rotula* de novo transcriptome-derived coding sequences (CDSs) and the recently sequenced *T. rotula* (strain FE80) (di Constanzo, et al., 2025) reference genome CDSs. Columns report Transcriptome-derived CDS hits to the genome (total number and percentage of CDSs with at least one BLASTp hit); Genome CDS hits to the transcriptome (total number and percentage of CDSs with at least one BLASTp hit); RBHs ("any": total number of reciprocal best hits, independent of thresholds and "filtered": retained after applying the specified sequence identity and coverage thresholds), shown as totals and as a percentage of all transcriptome-derived CDSs.

**mRNA read mapping stats:** number of reads mapped to *A. macleodii* CDSs, to *T. rotula* CDSs, and the total number of high-quality mRNA reads are reported per sample. Percentages indicate the fraction of total mRNA reads assigned to *A. macleodii*, *T. rotula*, or remaining unmapped. Mapping was performed at a nucleotide sequence identity threshold of 97% using BBMap.

**Supplementary Table S3 (Excel Worksheet)**

$Log_2$ fold-change values and adjusted p-values for *T. rotula* CDSs shown in Fig. 3.

**Supplementary Table S4 (Excel Workbook)**

Functional annotation of *T. rotula* genes included in the heatmap shown in Fig. 3, with the following tabs:

**T.rot_annot_Fig3** Columns summarize assigned biological processes or pathways and annotation results from multiple databases, including NCBI nr, UniProt, COG, KEGG, PFAM, and Gene Ontology (GO) terms. The last set of columns ("*T. rotula* genome") reports the best BLAST hits, percent identity, and KEGG/functional annotations based on the *T. rotula* FE80 reference genome (di Constanzo et al., 2025).

**T.rot_blast_Fig3** Results of blasts performed on four *T. rotula* genes included in the heatmap shown in **Fig. 3,** queried against sequences from other organisms. Columns report the Query sequence ID and the source organism, along with percent identity (pident), e-value, query coverage (qcov), subject coverage (scov), and gene product information (GPI).

**Supplementary Table S5 (Excel Worksheet)**
$Log_2$ fold-change values and adjusted p-values for *A. macleodii* CDSs shown in Fig. 4 Annotation (COG category and description, KEGG ortholog identifiers) is also reported.

**Supplementary Table S6 (Excel Worksheet)**
$Log_2$ fold-change values and adjusted p-values for *A. macleodii* CDSs additionally discussed in the text. Annotation (COG category and description, KEGG ortholog identifiers and, where applicable, CAZyme classification) is also reported.

**Supplementary movies**

**Movie 1.** Time-lapse light microscopy video of four *T. rotula* CCMP 3096 cells undergoing the morphological change, in co-culture with *A. macleodii* 27126. Time since the start of co-culture top left, hours:minutes. Scale bar, 20 µm.

**Movie 2.** Time-lapse light microscopy video of three *T. rotula* CCMP 3096 cells undergoing the morphological change, in co-culture with *A. macleodii* 27126. Video shows the formation of bacterial biofilm. Time since the start of co-culture top left, hours:minutes. Scale bar, 20 µm.

**Movie 3.** Time-lapse light microscopy video of *A. macleodii* 27126 chemotaxing toward a lysed *T. rotula* CCMP 3096 cell. The movie is displayed at a reduced playback speed for clarity; time since the start of the recording is shown at the top left (in seconds). Scale bar, 50 µm.

### Supplementary Discussion

#### Evidence that sexual reproduction is not induced in *T. rotula*

Despite the morphological similarity of the protoplasts with diatom sexual auxospores, sexual reproduction is not induced by *A. macleodii*. We tested whether diatom protoplasts were formed through an aborted sexual reproduction cycle by comparing *T. rotula*'s transcriptome with marker genes of diatom sexual reproduction[1–5]. We identified four homologous genes of the minichromosome maintenance family (*Mcm2*, *Mcm3*, *Mcm6* and *Mcm7*), which play a role in eukaryotic DNA replication during both mitosis and meiosis[6], as well as homologs of meiosis-specific genes that are expressed only during sexual reproduction (*Spo11, Msh5, Sig1, Sig2* and *Sig3*)[5,7,8]. These markers were either not differentially expressed in co-culture or were strongly downregulated (Fig. 3b; Supplementary Table S3), suggesting that sexual reproduction is not induced by the bacteria and that the observed protoplasts are neither oogonia nor auxospores.

#### Expression of genes related to nitric oxide signaling in *T. rotula*

We found no clear evidence for nitric oxide (NO) signalling, a key regulator of PCD in other algae[9] (Fig. 3b; Supplementary Table S3). Homologs of the NO-production enzymes *TpNoa*[9] and putative nitric oxide synthases (*Nos*)[10] showed only modest upregulation (max log2FC of 1.3 at 30 h), which is not consistent with strong production of NO to mediate the process. Similarly, nitrate and nitrite reductases (*Nar, Nir*) and associated transporters (*Ntrans*), capable of reductive NO production[11,12], were very strongly downregulated (log2fold changes < -7 at 30 h).

**Expression of carbohydrate metabolism genes in *A. macleodii***

*A. macleodii* utilized carbohydrates sparingly in co-culture, as evidenced by overall weak expression of carbohydrate-active enzymes (CAZymes), with a limited number showing sequential activation. Whilst 102 CAZymes were identified in the transcriptome, only a small number of these genes were significantly upregulated during co-culture with *T. rotula* (Supplementary Table S6). One exception was the strong and consistent upregulation of 5-keto-4-deoxyuronate isomerase (*KduI*), involved in the later stages of pectin degradation (Fig. 4b; Supplementary Table S5, log2FC > 2.9 at all time points). However, other genes from the pectin polysaccharide utilization locus (PUL) were not upregulated in co-culture, including *Pme* and *Pel* that act upstream in pectin degradation to produce the substrates of *KduI*. From Phase 2, we observed upregulation of CAZymes and transporters that indicate *A. macleodii* begins to utilise galactose and xylose-rich EPS polysaccharides, alpha-glucans and sucrose, an abundant metabolic intermediate in phytoplankton cells. In the third phase, particularly after 72 h, genes from the laminarin PULs[13] were upregulated (log2FC of all genes > 1.7). Laminarin is the primary storage carbohydrate in diatoms and could therefore be released when *T. rotula* cells lyse.

**Upregulation of siderophore related genes in *A. macleodii***

*A. macleodii* increased investment in iron acquisition in Phase 1, as shown by the upregulation of genes responsible for biosynthesis of the siderophore petrobactin (*Luc*) and two siderophore specific TonB-dependent transporters (*TbdT*)[14]. A number of other neighboring iron-responsive genes[14] (*SapC, PepS, Duf* (i) and *Duf* (ii)), were strongly induced in Phase 1, including an oxidoreductase (*FeO*) and tryptophan halogenase (*THal*) (Fig. 4b). The upregulation of *Dps*, which sequesters excess iron through a bacterioferritin domain to prevent oxidative damage, is indicative of elevated intracellular iron levels. Increased iron availability likely supported higher demand for heme cofactors, as indicated by the upregulation of protoheme IX farnesyltransferase, *CtaB* (log2FC 5.7 at 24 h). Correspondingly, genes encoding heme-containing components of the respiratory electron transport chain were upregulated (Fig. 4b), including cytochrome c oxidase subunits (*CoxI–III*) and assembly factor (*CtaG*). Heme cofactors also support the activity of the nitrate assimilation enzymes *NirB-NasA-CobA*[15,16], which were likewise upregulated.

**Potential for methanol oxidation by *A. macleodii***

Genes for the biosynthesis of pyrroloquinoline quinone (PQQ) were induced from 5 and 10 h in *A. macleodii* (Fig. 4b), along with two periplasmic PQQ-dependent alcohol dehydrogenases (*Adh*), all reaching log2FC > 3 at 24 h. Two additional genes, encoded in the same neighbourhood as *Adh (ii)*, were also very strongly upregulated (log2FC >6 24-72 h); cytochrome c-550 electron acceptor *PedF*, which can serve as an electron acceptor for PQQ-dependent *Adh*, and an NAD-dependent aldehyde dehydrogenase (*Aldh* (iii)), which could catalyze oxidation of the resulting aldehyde to acid. Whilst *A. macleodii* also encodes enzymes for the downstream oxidation of formaldehyde via THF- and GSH-link pathways, these were not upregulated in co-culture. However, we detected modest upregulation of *FdhA*, a subunit of formate dehydrogenase (log2FC > 1 all timepoints), capable of oxidising formate to $CO_2$. This suggests that *A. macleodii* may have the capacity for complete oxidation of methanol to $CO_2$, though further biochemical validation is needed to determine whether methanol or other alcohols serve as the principal substrate. *A. macleodii* strain ATCC 27126 is reportedly unable to grow on methanol as a sole carbon source, consistent with the absence of a complete serine cycle[13,16]. In addition, we observed strong upregulation of several molybdo–xanthine oxidoreductase (XOR)–family genes ($log_2FC > 6$; Supplementary table S6), which catalyse oxidation of diverse organic compounds, again with limited substrate annotation.

Given the strong upregulation of PQQ biosynthesis, it is also relevant that PQQ has functions beyond metabolism, including interconversion of dioxygen-superoxide, acting as a potent antioxidant and signal regulator, for example in mitochondrial and peroxisomal biogenesis.
